# Systematic Analysis of the EXO70 Gene Family in Kiwifruit Species: Evolutionary Selection and Potential Functions in Plant Immunity

**DOI:** 10.1101/2025.10.28.684437

**Authors:** Wei Cui, Cecilia Deng, Minsoo Yoon, Viktor Žárský, Erik Rikkerink

## Abstract

**Background:** Kiwifruit (*Actinidia* spp.) is a commercially and nutritionally valuable fruit crop that faces increasing challenges from pathogens, particularly *Pseudomonas syringae* pv. *actinidiae* (Psa). The exocyst complex, especially the EXO70 subunit, has been demonstrated to play a crucial role in vesicle trafficking and plant immune responses in model species such as *Arabidopsis thaliana*. However, the function and evolution of EXO70 genes in fruit crops remain largely unexplored.

**Results:** We conducted a comprehensive genome-wide analysis of the EXO70 gene family across five *Actinidia* species using *Arabidopsis* EXO70 sequences as queries. A total of 217 EXO70 genes (23 to 54 paralogues per species) were identified and classified into three subfamilies and nine clades (EXO70A-EXO70I), consistent with previous classifications in other plant taxa. Phylogenetic reconstruction and microsynteny analyzes revealed lineage- and genus-specific expansions of EXO70C members, as well as species-specific expansion events, particularly within the EXO70E and EXO70H clades. To investigate potential immune functions, yeast two-hybrid and *in planta* co-immunoprecipitation assays confirmed that kiwifruit EXO70B1 physically interacts with the immune hub protein kiwifruit RIN4_1. These interactions support conservation of the EXO70-RIN4 module in plant immunity.

**Conclusions:** This study provides the first comprehensive characterization of the EXO70 gene family across multiple kiwifruit species and uncovers candidate genes potentially involved in plant immunity through RIN4-mediated defense signaling. Notably, the expansion and diversification of the EXO70E and EXO70H clades suggest possible adaptive evolution within these clades and neofunctionalization. These findings provide valuable genomic resources and novel insights into the evolution of vesicle trafficking components in fruit crops, laying the groundwork for future efforts to enhance disease resistance in kiwifruit through breeding or biotechnological approaches.

## 1. Introduction

Kiwifruit (*Actinidia* spp.) is cultivated globally for its nutritional value and commercial importance. However, production is increasingly threatened by diverse biotic stresses, including bacterial and fungal pathogens and pests. Among these, bacterial canker caused by *Pseudomonas syringae* pv. *actinidiae* (Psa) is a particularly severe disease that has had significant economic impact on kiwifruit industries worldwide [1]. Developing effective and durable resistance strategies requires a deeper understanding of the molecular components underpinning host immune responses.

In plants, targeted vesicle trafficking is essential for cell growth, signaling, and defense. This process is mediated by the exocyst complex, a conserved octameric complex comprising SEC3, SEC5, SEC6, SEC8, SEC10, SEC15, EXO70, and EXO84 subunits. Of these, EXO70 has undergone remarkable expansion in plants [2,3]. While fungi and animals typically carry a single EXO70 gene, *Arabidopsis thaliana* encodes 23 EXO70 genes, which are grouped into eight major clades (EXO70A to EXO70H) and some species have an additional ninth clade (EXO70I). These genes encode proteins with diverse functions, including cell wall remodeling, polar growth, and immune signaling [4].

Emerging evidence highlights a role for EXO70 proteins in plant immunity, often through interactions with immune regulators and/or pathogen effectors. For example, in rice, EXO70F2 acts as a decoy for the fungal effector AvrPii and cooperates with the resistance protein Pii [5,6]. In *Arabidopsis*, EXO70B1 and EXO70B2 contribute to FLS2 receptor recycling, stomatal regulation and interact with the immune regulator RIN4 (RPM1-Interacting Protein 4) [7–9]. RIN4, an intrinsically disordered protein is a central hub targeted by several bacterial effectors [10], is also known to mediate immune responses through its interaction with multiple resistance (*R*) proteins [11–16]. Several EXO70B and EXO70E isoforms physically interact with the disordered immune hub protein RIN4 [17–19]. Moreover – growing evidence indicates that members of EXO70.2 subfamily, including RIN4 interactors EXO70B and EXO70E, are integrated into the autophagy and unconventional protein secretion related processes also in defence [20–24].

Despite their importance, EXO70 genes remain poorly characterized in fruit crops, including kiwifruit. The recent availability of genome assemblies for several *Actinidia* species provides an opportunity to investigate phylogenetically the EXO70 gene family across the genus and to assess their potential roles in plant defense.

In this study, we aimed to (**i**) identify and classify EXO70 genes in five *Actinidia* species; (**ii**) investigate their evolutionary relationships and potential signatures of diversifying selection; and (**iii**) explore interactions between EXO70 clades and RIN4 using yeast two-hybrid and co-immunoprecipitation assays. By integrating phylogenomic, synteny, and functional analyzes, we provide new insights into the evolution and immune function of EXO70 proteins in this non-model fruit crop. Our findings offer a foundation for future research and potential breeding strategies to enhance disease resistance in kiwifruit.

## 2. Results

### 2.1. Genome-Wide Identification and Classification of EXO70 Genes in Kiwifruit

We identified a total of 217 EXO70 genes from six genomes across five *Actinidia* species -*Actinidia chinensis* var. *chinensis* (*A. chinensis* var. *chinensis*, Red5 reference genome and Hongyang), *A. melanandra*, *A. arguta*, *A. polygama* and *A. eriantha* - through homology-based searches using *Arabidopsis thaliana* ‘Columbia’ EXO70 sequences and *Solanum lycopersicum* EXO70I as references. These genes were classified into three subfamilies (EXO70.1, EXO70.2, and EXO70.3) and nine major clades (EXO70A to EXO70I), consistent with established classifications in model plants and other angiosperms [25].

Phylogenetic analysis revealed that all nine clades were represented in *Actinidia*, including symbioses regulating EXO70I, which is absent in *Arabidopsis* [26]. Gene counts varied across species, reflecting differences in genome assembly quality, ploidy levels, and annotation strategies. Notably, *A. melanandra* harbored the largest number of EXO70 genes (n = 54), likely due to the inclusion of uncollapsed haplotypes in its draft genome. A summary of EXO70 gene counts per species is provided in **Table 1** and the phylogeny of EXO70 clades in the Red5 - *Actinidia* reference genome is presented **Figure 1**.

**Figure 1:**
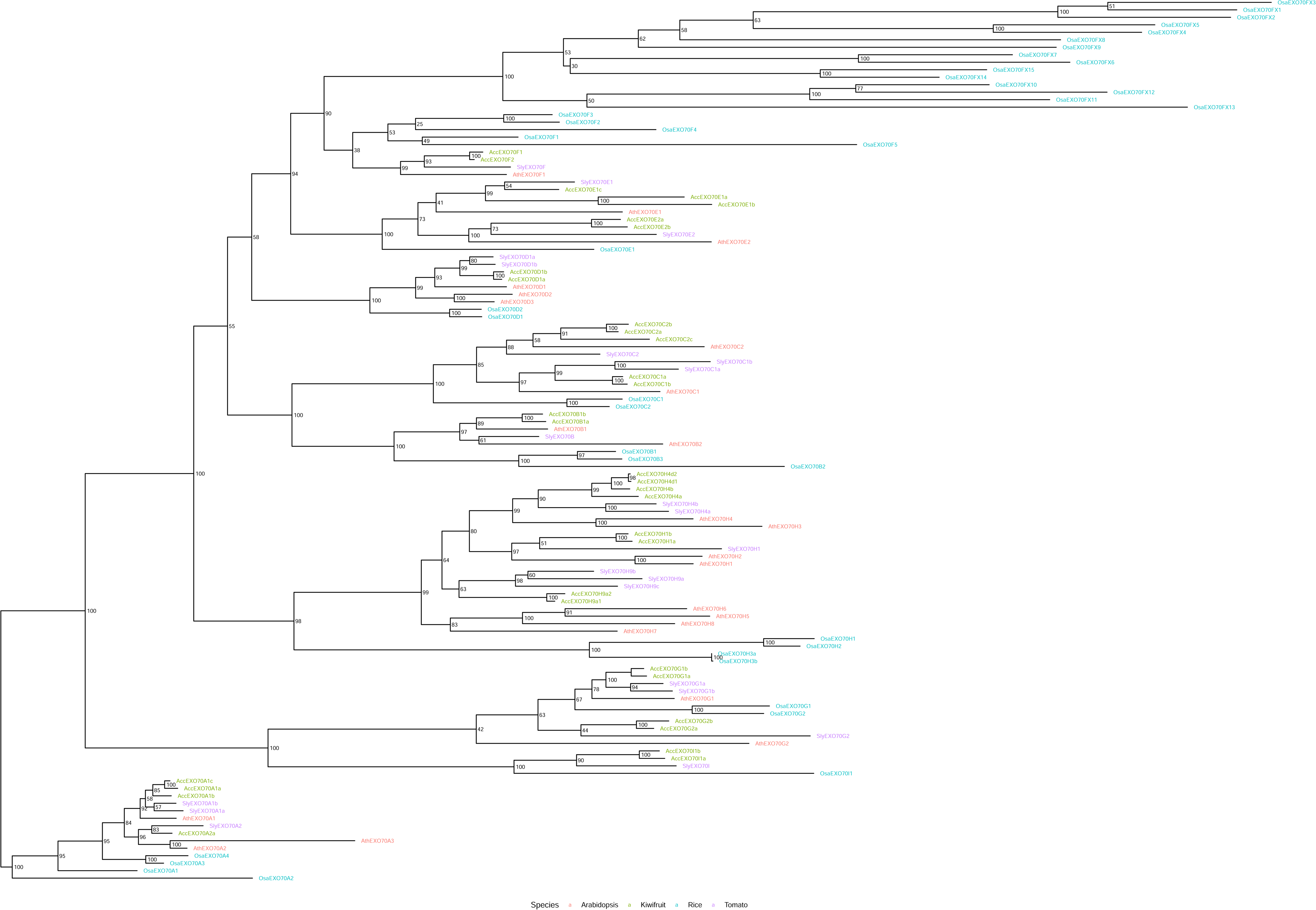
Phylogenetic tree of MAFFT v7.307 protein sequence alignment using RAxML v8.2.11. The tree shows the relationships among EXO70 proteins from multiple *Actinidia* species, *Arabidopsis thaliana*, tomato and rice. The nine major clades (A–I), corresponding to the EXO70A-EXO70I subfamilies. Proteins are color coded according to species of origin (kiwifruit green; *Arabidopsis* red; tomato purple; rice light blue). Tree tips are labeled with individual gene identifiers from each species.

**Table 1.** EXO70 genes identified in each subfamily across six *Actinidia* genomes, *Arabidopsis*, tomato and rice.

| Species/genotype | *<sup>1</sup>Three letters | Total | A | B | C | D | E | F | G | H | I |
| --- | --- | --- | --- | --- | --- | --- | --- | --- | --- | --- | --- |
| *Actinidia chinensis* var. *chinensis* Red5 | Acc | 34 | 4 | 2 | 5 | 2 | 5 | 2 | 4 | 8 | 2 |
| *Actinidia chinensis* var. *chinensis* Hongyang | Ach | 23 | 4 | 2 | 5 | 1 | 2 | 1 | 2 | 4 | 2 |
| *Actinidia melanandra* | Ame | *<sup>2</sup>54 | 10 | 5 | 6 | 7 | 5 | 2 | 2 | 16 | 1 |
| *Actinidia arguta* | Aar | 36 | 4 | 2 | 5 | 2 | 5 | 3 | 4 | 9 | 2 |
| *Actinidia polygama* | Apo | 36 | 5 | 2 | 5 | 2 | 5 | 2 | 4 | 9 | 2 |
| *Actinidia eriantha* | Aer | 34 | 4 | 2 | 5 | 2 | 7 | 1 | 3 | 8 | 2 |
| Total proteins in *Actinidia* clade specific trees |  | **217** | **31** | **15** | **31** | **16** | **29** | **11** | **19** | **54** | **11** |
| **Model Plant Genomes** |  |  |  |  |  |  |  |  |  |  |  |
| *Arabidopsis thaliana* Col | Ath | 23 | 3 | 2 | 2 | 3 | 2 | 1 | 2 | 8 | *<sup>3</sup>0 |
| *Solanum lycopersicum* cv. Heinz 1706 | Sly | 22 | 3 | 1 | 3 | 2 | 2 | 1 | 3 | 6 | *<sup>4</sup>1 |
| *Oryza sativa* cv. Nipponbare | Osa | 39 | 4 | 3 | 2 | 2 | 1 | *<sup>5</sup>20 | 2 | 4 | 1 |

Summary of the number of genes identified in each of the nine EXO70 subfamilies in six *Actinidia* genomes and the genomes of model plants. \*<sup>1</sup>Three letter abbreviation code that distinguishes between genes derived from different genomes used to name all genes/proteins; \*<sup>2</sup>The *A. melanandra* genome used in this analysis is a draft genome that likely includes multiple copies and haplotypes and probably accounts for the larger number of candidate proteins; \*<sup>3</sup>The EXO70I clade is not present in Brassicaceae such as *A. thaliana*; \*<sup>4</sup>SlyEXO70I has been used as a reference gene in the EXO70I clade specific trees whereas *Arabidopsis* genes serve as reference genes in the subsequent clade specific trees; \*<sup>5</sup>This includes five genes in the EXO70F clade and 15 in the monocot specific EXO70Fx subclade.

Locus names were originally assigned after alignments and largely remained consistent through phylogenetic reconstruction. In rare instances where the gene nomenclature had to be adjusted to fit phylogenetic placements in the clade specific trees, we documented the original names in **Table 1**. These nomenclature adjustments aid clarity when referring to specific subclades or branches.

### 2.2. Selection analysis of EXO70 Clades

After classifying the 217 EXO70 proteins identified from six *Actinidia* genomes into nine clades we constructed clade specific phylogenetic trees containing the proteins from all the *Actinidia* species represented in the dataset (**Figure 2** for EXO70E and EXO70H clades, supplementary **Figures S1-S7** for all other clades). One apparent feature of all genus *Actinidia* genomes is evolutionary multiplication of “non-canonical” EXO70C subfamily. The resulting trees were then inspected to identify potential signs of selection. The indicators of selection identified included: unusually long branch lengths of individual proteins, frequent gene loss/gain events in particular subclades and incongruous patterns of relationships between EXO70 loci not predicted by phylogenetic relationship previously established between the kiwifruit species in this analysis [27]. Presence/ absence analyzes of loci in gene sets processed by different methods can be compromised by a lack of consistency between annotations. To compensate for inconsistencies in annotation and either support or refute these presence/ absence polymorphisms, we investigated potential gene-loss scenarios within the EXO70 clades by tBlastN analysis of the *Actinidia* genomes. These searches used available *Actinidia* genome assemblies in the NCBI WGS database. This found evidence for genome sequence matching most of the unannotated gene loci. We also investigated unusually long tree branches by tBlastN searches of relevant kiwifruit genomes. In many cases the variation in sequence that gave rise to these long branches was due to either incomplete or incorrect gene annotations, or splicing variants. The results of these analyzes are provided below the relevant supplementary figures (**Figures S1-S7**). There were instances where no matching genome sequence was identified or variants that had sustained significant modifications at the protein level were identified and that explained the long branch lengths. At the conclusion of this preliminary screen and supporting investigations we identified subclades with significant intrageneric signs of selection within the EXO70E and EXO70H clades.

**Figure 2:**
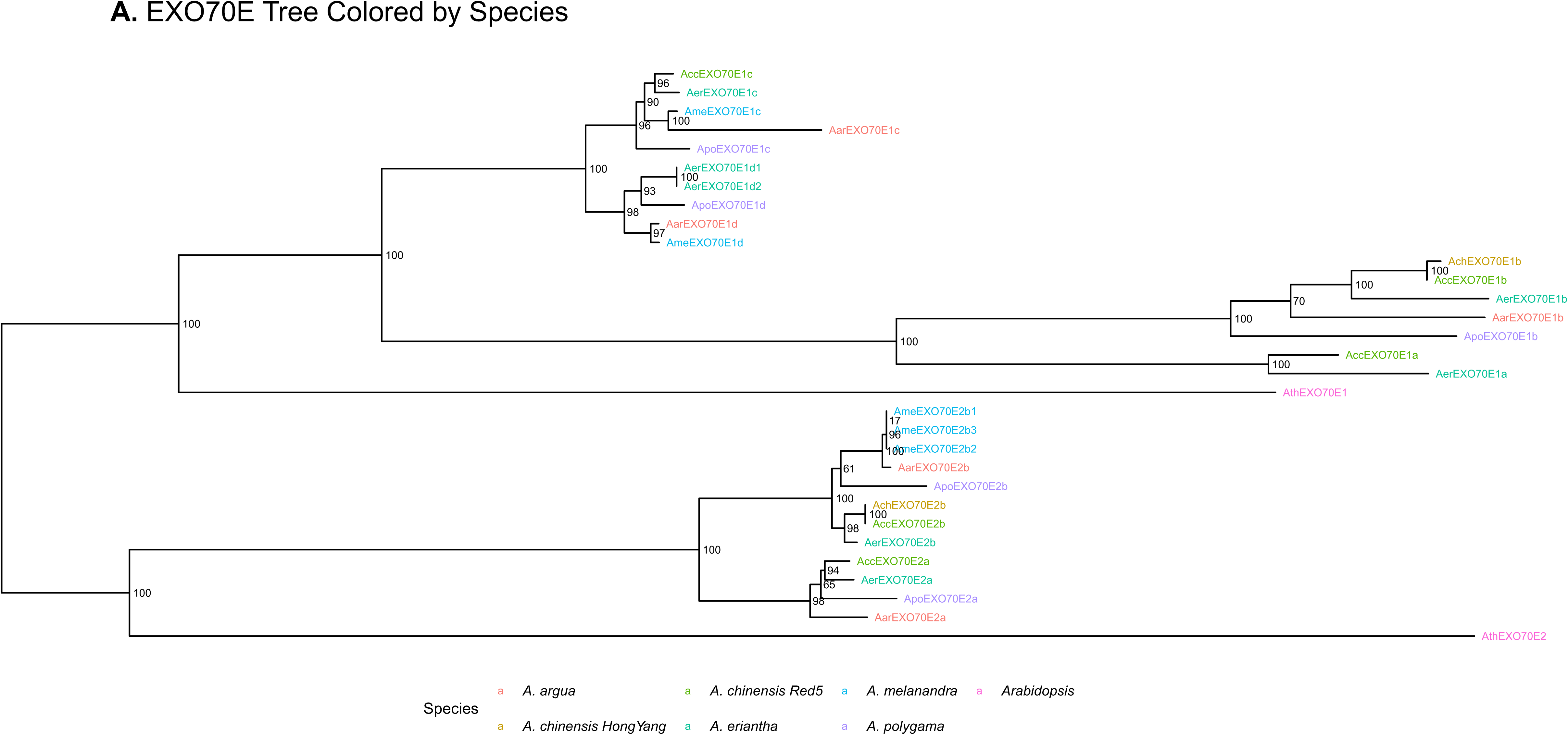
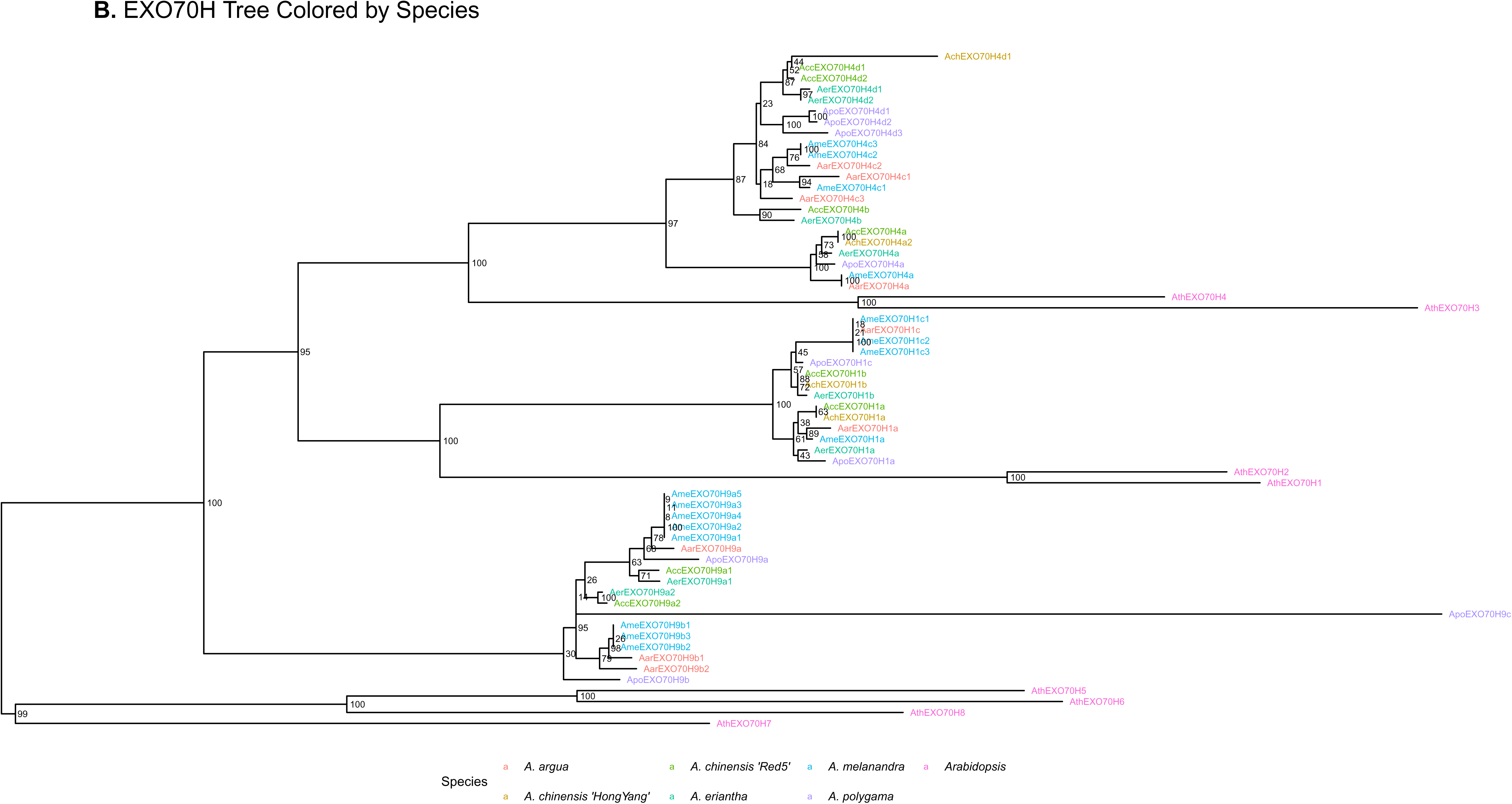
EXO70 clade specific phylogenetic trees. **A**) Clade specific tree of EXO70E clade proteins with genome of origin of the proteins identified by color (*A. arguta*- red; *A. chinensis* var. *chinensis* Red5 -green and Hongyang -olive; *A. eriantha* -turquoise; *A. melanandra* -light blue; *A.polygama* purple; and *A. thaliana* -pink). **B**) Clade specific tree of EXO70H clade proteins using the same identifying colors as above.

### 2.3. Divergence, Amplification and Gene Loss in the EXO70E1 Subclade

Some of the genes identified as potentially missing from analysis of the EXO70E phylogenetic tree were in fact present but unannotated. For example, a locus with 99% identity to *Ach*EXO70E1c was identified in *A. chinensis* Hongyang, and a match to *Acc*EXO70E2a was found in *A. melanandra*, despite both being absent from predicted gene models. In contrast, no ortholog of EXO70E1a was detected in *A. polygama*, suggesting likely true gene loss in this species. EXO70E1b was also truncated in *A. polygama* and both EXO70E1a and EXO70E1b were predicted as truncated or divergent in *A. melanandra* and *A. arguta*. In contrast, EXO70E1c was conserved across all species and served as a stable phylogenetic reference in the EXO70E1 subclade (**Table S1**). Comparison of the translation alignment between EXO70E1a and EXO70E1b indicated that the EXO70E1a locus is likely to have sustained a large in frame deletion in the middle of the EXO70E1 coding region. Remarkably this deletion encompassed the two alpha helices identified as the most highly conserved in a ConSurf analysis of the *Ath*EXO70B1 protein (**Figure S8**) [28,29].

The variant of the EXO70E1c locus in *A. arguta* (*Aar*EXO70E1c) displayed long branch lengths and low identity to orthologs, particularly when compared with the locus from its near neighbour *A. melanandra*. A follow-up tBlastN search identified these differences were probably due to annotation errors with incorrect start and ending sequences responsible for most of the differences. An alternative protein reading frame 99.6% identical to *Ame*EXO70E1c was present on chromosome 27 in *A. arguta*.

Our sequence tBlastN searches also identified a new EXO70E1 locus in the Red5 genome, located on chromosome 5 within a region syntenic to chromosome 27 (Genomicus syntenic block 63, **Figure S9**) that contains the *Acc*EXO70E1c locus. When this new Red5 reference gene was used to search other *Actinidia* genomes it was invariably also found in these genomes on chromosome 5. This new locus was named EXO70E1d. The EXO70E1a and EXO70E1b loci are closely linked to each other (between just 3.5 kb and 26 kb apart) in all the *Actinidia* genomes where this could be analyzed. Given the locations of the EXO70E1a and EXO70E1b loci, this supports the occurrence of two sequential duplications within the EXO70E1 subclade to give rise to the four members of this clade (**Figure 2A**).

Within the EXO70E clade, members of the EXO70E1 subclade showed sequence divergence, amplification, gene truncation, and species-specific absence (**Table S2**). Given the evidence for selection in this clade, a more comprehensive test for evidence of episodic selection in the EXO70E1 clade was undertaken using the Datamonkey 2.0 web site [30,31]. This analysis identified significant evidence for selection having operated on the EXO70E1a and EXO70E1b branches, and within the *Ach*EXO70E1b and *Acg*EXO70E1a (*<u>A</u>. <u>c</u>hinensis* var. *chinensis* <u>G</u>uimi No. 2) proteins based on reasonable likelihood ratios and empirical Bayes factors (**Figure S10** and **Table S2**).

### 2.4. Amplification in the EXO70H4 Subclade

Within the EXO70H clade an *Apo*EXO70H9c allele was only found only in *A. polygama*, this is likely a duplicated, degenerating copy of *Apo*EXO70H9b based on tBlastN analysis of the Hokkaido genome. The likely sequence degeneration also accounts for the very long branch length. Incongruent branching can also be seen for *Apo*EXO70H9b but this has weak support (30%), making the evidence for selection weaker.

In contrast, the EXO70H4 subclade showed compelling evidence for recent species-specific gene amplification. Several paralogous copies were detected in *A. chinensis* and *A. arguta*, with phylogenetic placements that suggest independent amplification events. These patterns suggest that EXO70H4 loci are evolving rapidly, likely under selective pressures unique to specific *Actinidia* lineages. The combination of high sequence variation and phylogenetic incongruence suggests recent and independent expansion events within this subclade (**Figure 2B**).

### 2.5. Interaction of EXO70 Proteins with RIN4

Previous *Arabidopsis* research has linked several EXO70 proteins to plant defense *via* their interaction with the disordered immune regulator RIN4 [17–19]. To explore the functional relevance of EXO70 proteins in *Actinidia*, we tested for selected protein-protein interactions with kiwifruit RIN4. *Acc*EXO70B1 and *Acc*EXO70E2 exhibited strong interactions with *Acc*RIN4_1 in yeast two-hybrid assays (**Figure 3A**). This interaction between *Acc*EXO70B1 and *Acc*RIN4_1 was subsequently validated *in planta* using co-immunoprecipitation assays in *Nicotiana benthamiana* (**Figure 3B**). Notably, not all EXO70 proteins showed interaction with RIN4. For instance, *Acc*EXO70A1, did not exhibit binding activity, suggesting that EXO70-RIN4 associations are specific to certain clades. The observed interactions in kiwifruit mirror those reported in *Arabidopsis* [17,32,33], supporting a conserved role for at least some of the clade-specific EXO70-RIN4 complexes in plant immunity.

**Figure 3:**
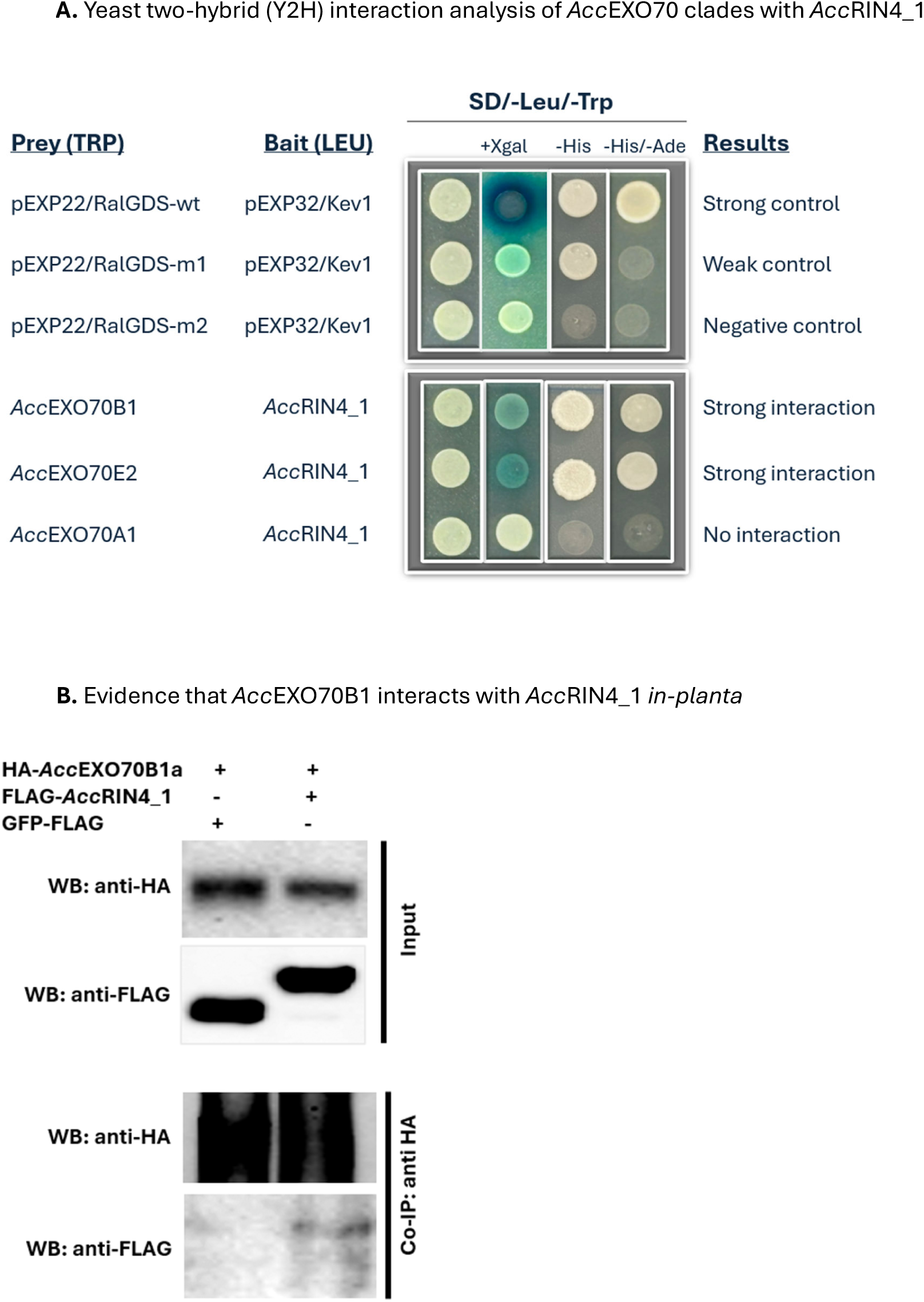
Interaction of selected kiwifruit EXO70 proteins with RIN4 in yeast and *in planta*. **A).** The *Saccharomyces cerevisiae* strain MaV203 was co-transformed with *Acc*RIN4_1 bait (pDEST32) and prey constructs of different *Acc*EXO70 isoforms (pDEST22). Transformants were selected on SD/-Leu/-Trp medium and tested for interaction by growth on SD/-Leu/-Trp/-His/-Ade and by LacZ reporter activation on SD/-Leu/-Trp plates supplemented with X-gal. Blue coloration indicated a positive interaction. Strong interactions were detected between *Acc*EXO70B1a and *Acc*RIN4_1 and between *Acc*EXO70E2a and *Acc*RIN4_1, whereas no interaction was observed for *Acc*EXO70A1. Positive and negative control pairs (strong, weak and non-interacting) were included to validate the assay. **B).** Evidence that kiwifruit EXO70B1 (*Acc*EXO70B1a) interacts with kiwifruit RIN4_1 (*Acc*RIN4_1), demonstrated with *in planta* co-immunoprecipitation (co-IP) analysis. For co-immunoprecipitation of *Acc*EXO70B1a and *Acc*RIN4_1. *Nicotiana benthamiana* leaves were syringe-injected using an *Agrobacterium* mixture harboring HA:*Acc*EXO70B1a and FLAG:*Acc*RIN4_1 in equal amounts. Protein extracts were immuno-precipitated using anti-HA antibody. Western blots prepared from the precipitated proteins were probed with anti-HA and anti-FLAG antibodies as indicated.

Overall, these results underscore the evolutionary and functional diversification of the EXO70 family in kiwifruit and highlight specific EXO70 members as promising candidates for further study in the context of understanding both defense signaling and disease/ pest susceptibility.

## 3. Discussion

This study presents the first comprehensive analysis of the EXO70 gene family in kiwifruit (*Actinidia* spp.), uncovering its evolutionary diversification and potential role in plant immunity. By integrating comparative genomics, phylogenetic reconstruction, and functional assays, we identified clade-specific expansions and signatures of diversifying selection in EXO70 genes, particularly within the EXO70E and EXO70H clades. These findings suggest that certain EXO70 clades have undergone neofunctionalization, likely in response to selective pressures imposed by pathogens and pests.

### 3.1. Evolutionary Expansion and Contraction of the EXO70 Gene Family in Kiwifruit

We identified 217 EXO70 genes across five *Actinidia* species, with notable variability in gene counts among species. This variation reflects differences in genome assembly quality and ploidy, but also included genuine lineage-specific expansions, gene loss and gene inactivation. The largest number of EXO70 loci was observed in *A. melanandra*, largely due to haplotype redundancy in its draft genome. Despite these technical caveats, phylogenetic and syntenic analyzes supported a core classification into nine conserved clades (EXO70A– EXO70I), consistent with previous studies in *Arabidopsis* and other angiosperms.

*Actinidia* underwent a duplication event at or near the origin of the genus (hereafter named Ac-α) which has resulted in most genes being present as duplicates in syntenic chromosome locations. The EXO70E and EXO70H clades showed expansion beyond Ac-α duplication across *Actinidia* genomes. EXO70E1 exhibited strong evidence for diversifying selection, including frequent gene loss/ inactivation, truncation, divergence, and likely non-functionalization. At least one of these signatures occurred in all the analyzed species, suggesting independent and possibly convergent evolutionary trajectories within this subclade. In contrast, EXO70E2 showed more conserved patterns, with only the two copies derived from Ac-α and limited evidence of recent diversification.

Similarly, the EXO70H4 subclade showed recent amplification events in *A. chinensis* and *A. arguta*. Amplification signatures within these clades are consistent with previous reports of EXO70 involvement in lineage-specific stress responses and suggest functional specialization that is most likely to be related to biotic stress adaptation. Given the important role of EXO70 proteins in a number of plant defense processes [5–9] and the expansion of this gene family it is plausible that certain EXO70 genes could end up being specifically targeted by particular insects or pathogens and thus act as “susceptibility” loci with respect to these insects or pathogens. Thus absence or frequent inactivation of loci from a genome might be an indicator that suggests selection operating against such loci. Our analysis indicates that interpreting gene loss requires particular caution. Other than biological reasons, such losses may also have alternative explanations. Other possible explanations include incomplete genome assembly, insufficient transcriptomics data to support gene model prediction, and/ or less rigorous or inconsistent annotation processes between genomes. Although a few cases of complete gene loss appear to be real, most of the cases of genes entirely missing turned out to be due to one of these alternative reasons. A high frequency of gene inactivation is another pattern that forms equally convincing evidence for selection. In the case of the EXO70E1a sub-clade, the majority of *Actinidia* species and varieties showed inactivation of this particular sub-clade by frame shifts and/ or the evolution of premature stop codons. This inactivation strongly suggests this locus may be acting as a susceptibility locus against some pests and/or diseases in kiwifruit. This hypothesis is further supported by this branch showing the most significant signs of selection based on the aBSREL analysis. The branch with the second most likely signatures of selection based on our analysis was the closely related EXO70E1b locus. This locus occurs adjacent to the EXO70E1a locus based on the microsynteny analysis and these two loci show the greatest similarity to each other in alignments. A comparison between the alignment of the complete EXO70E1a and EXO70E1b loci indicates a large in-frame deletion occurred in the middle of the EXO70E1a proteins that interferes with the two most highly conserved α helices identified in the Consurf analysis and this region is also the site of interaction of the fungal effector AvtPii with *Osa*EXO70F2 [6,29]. Given that this region is apparently also targeted by effectors, it is tempting to speculate that this deletion could link more directly to a novel functional adaptation for EXO70E1a.

### 3.2. Functional Diversification and Immune Relevance

Our yeast two-hybrid assays confirmed that *Acc*EXO70B1 and *Acc*EXO70E2 physically interact with *Acc*RIN4_1, a key immune regulator. The interaction between *Acc*EXO70B1 and *Acc*RIN4_1 was further validated *in planta* using co-immunoprecipitation. These results mirror interactions observed in *Arabidopsis* and rice, extending the relevance of the EXO70– RIN4 module to a perennial fruit crop. The fact that not all EXO70 isoforms interact with RIN4 further supports the idea of functional specialization among EXO70 family members, as proposed previously [25,33].

The immune relevance of EXO70 clades is increasingly supported by functional studies in model plants. For example, EXO70B1 and B2 are known to regulate pattern recognition receptor (PRR) trafficking and participate in *R*-protein–mediated responses [7,9]. EXO70E and EXO70H members have also been implicated in immune signaling, sometimes acting as effector targets or decoys [5]. In rice, EXO70F2 interacts with the AvrPii effector and is guarded by the resistance protein Pii [6]. A member of the EXO70H clade in rice was recently shown to interact with the insect resistance protein BPH6 [34], and expression analyzes suggest EXO70H members are upregulated under biotic stress. These parallels highlight the likelihood that diversification in the *Actinidia* EXO70E1 and EXO70H4 clades is similarly driven by host–pathogen co-evolution.

### 3.3. Phylogenetic Insight into Selection and Clade Specialization

Our clade-specific phylogenetic reconstructions revealed that EXO70E and EXO70H clades are among the most rapidly expanding EXO70 lineages in *Actinidia* (along with EXO70C). The deep evolutionary divergence of the EXO70E1/E2 split/ absent in monocots but present in basal eudicots, suggests early subfunctionalization in dicots followed by lineage-specific diversification. Whether this diversification is directly linked to its ability to interact with RIN4 is currently unknown. In contrast, the other EXO70 proteins showing interaction with RIN4, from the EXO70B clade remain relatively conserved in kiwifruit, with only the two loci derived from the genome wide Ac-α duplication identified.

Interestingly, the EXO70B2 clade appears restricted to Brassicaceae, where it cooperates with EXO70B1 in the regulation of FLS2-mediated immunity [9]. These data contrast with findings in legumes, where the EXO70J clade (proposed to be renamed as EXO70BX given it is related to EXO70B) underwent massive expansion, possibly to support nodulation and nitrogen fixation [35–37]. The lack of dramatic amplification within the EXO70B1 clade in *Actinidia* (and other Angiosperm species) suggests either functional conservation or the absence of selection forces similar to the pressures operating in legumes.

### 3.4. Comparative Genomics Enables Detection of Evolutionary Signals

Our multi-species comparative approach was essential for identifying signatures of selection. Initially, no strong diversification patterns were obvious in the reference genome of *A. chinensis* Red5. However, expanding the analysis to five additional *Actinidia* genomes revealed clear signals of selection and diversification in the EXO70E1 and EXO70H4 subclades. This highlights the importance of deep phylogenomic sampling to detect rapid adaptive gene family evolution in cases where the signatures may be genus or species specific.

While many EXO70 clades followed expected species relationships, deviations in EXO70E1 and EXO70H4 trees - such as unusually long branches or incongruent clustering - suggest adaptive evolution and functional divergence. Although not all of these patterns survived closer scrutiny (see below) at least some of these patterns were supported by conserved microsynteny, BLAST validation, and further gene model curation, increasing the likelihood of discovering future biological reasons for our findings.

### 3.5. Caveats in Interpreting Selection Signatures

We acknowledge that signs of diversification must be interpreted cautiously. Differences in genome quality and annotation can mimic evolutionary signals. For example, long branches can result from mispredicted open reading frames, and inflated gene counts may arise from comparing genomes with different strategies for conveying haplotype redundancy. We addressed these issues by curating sequences through tBLASTn verification and by excluding incomplete or dubious gene models from phylogenetic analyzes. Our analysis thus reveals a pathway by which the growing body of genome data can be used for similar analyzes, even if it has not all been processed initially in the same way or with the same degree of rigour in annotation. Despite the limitations, the use of multiple *Actinidia* genomes allowed robust filtering of false positives and revealed broadly consistent patterns across species. Subclades EXO70E1 and EXO70H4 emerged as the most promising candidates for further functional investigation.

### 3.6. Implications and Future Directions

This work lays a foundation for future functional studies of EXO70-mediated immunity in fruit crops. Our findings suggest that EXO70E1 and EXO70H4 clades may play specialized roles in *Actinidia* defense, possibly through interactions with immune signaling hubs like RIN4. Given their lineage-specific amplification and divergence, these two clades represent promising targets for accelerating disease resistance breeding and/or genome editing.

Future research should integrate expression profiling, pathogen challenge assays, and other investigations to understand the nature of the role of this gene family in conferring susceptibility and/ or resistance in kiwifruit. Clarifying whether these EXO70E1 and EXO70H4 proteins are involved in interactions that activate effector-triggered immunity would deepen our understanding of how vesicle trafficking intersects with immune signaling in perennial fruit crops.

## 4. Materials and Methods

### 4.1. Genome Resources and Gene Identification

*A. chinensis* Hongyang v3 [38] was downloaded from NCBI (PRJNA549770). Genomes and annotations for *A. arguta* Ruanzao W1 [39], *A. eriantha* Maohua W1 [40] and *A. polygama* Gezao W1 were downloaded from the Kiwifruit PanGenome Database [41]. Peptide sequences for genes annotated as ‘EXO70’ in each assembly were extracted as candidate EXO70 genes. *A. melanandra* genome assembly ME02_01 was downloaded from NCBI (PRJNA1080037) and indexed with miniport [42]. EXO70 peptide sequences from *Arabidopsis* (TAIR10) and Hongyang v3’ were used as query and aligned to ME02_01 genome with miniport to predict EXO70 genes in ME02_01. The peptide sequences were then extracted from the genome using AGAT v1.0.0 [43].

Genome abbreviations were attached to the start of gene IDs of candidate EXO70 genes from each *Actinidia* species. They were combined to serve as a protein database. Hidden Markov Model (HMM) profiles of the EXO70 domain (PF03081) were used to search the databases using HMMER3.0 [44–47]. Identified genes were subsequently verified by conserved domain analyzes using SMART [48,49] and Pfam [50] databases, yielding a clean set of EXO70 protein sequences for the selected kiwifruit genomes. EXO70 proteins were classified into clades and subclades based on preliminary alignments with EXO70 proteins from *A. thaliana*, tomato, and rice using default settings in Geneious Prime 2025.0.2 (https://www.geneious.com). These preliminary alignments were used to identify *Actinidia* proteins covering less than 50% of the consensus sequence that were removed from the dataset to yield the final set of kiwifruit EXO70 proteins. The annotations of genes with particularly long branches in phylogenetic trees (below) were re-examined to identify if these long branches are likely to be due to annotation errors.

### 4.2. Phylogenetic and Synteny Analysis of EXO70 Clades

A naming system consistent with the *Arabidopsis* EXO70A–EXO70H clade nomenclature was adopted to support comparative analysis and discussion. Clades were denoted by capital letters (EXO70A–EXO70I), while subclades were distinguished by adding numbers (e.g., EXO70H1–H8) to reflect their counterparts in *Arabidopsis* where possible. Novel subclades not found in *Arabidopsis* were assigned new numbers (e.g., EXO70H9). Additional members within subclades were further differentiated using lowercase letters and numbers (e.g., EXO70E1c1) to aid in the identification and analysis of larger subclades. EXO70I proteins, absent in *Arabidopsis*, were identified and named based on homology with tomato (*Solanum lycopersicum* cv. Heinz) EXO70I proteins.

Protein sequences from each clade were aligned using MAFFT v7.307 and trimmed with trimAI v1.5.0. Phylogenetic trees were generated using RAxML v8.2.11 under the PROTGAMMA model with 1,000 bootstrap replicates, and visualized with Dendroscope v3. Final annotated trees were produced using the ggtree and ggplot2 packages in R v4.4.2.

Gene naming was further supported by microsynteny analysis using GenomicusPlants v49.01 (https://www.genomicus.bio.ens.psl.eu/genomicus-plants-49.01/) and EnsemblPlants gene trees. Microsynteny aided in the classification of genes not identified in previous analyzes such as EXO70E1d found in a region homoeologous to the EXO70E1c locus. Microsynteny supported subclade assignments across dicot species and was useful in identifying “sister” loci in *Actinidia* species linked to the Ac-α whole genome duplication that likely predates the evolution of the genus. Given that only the Red5 genes from *A. chinensis* were represented in EnsemblPlants, these genes served as reference anchors for cross-species orthology assignments in *Actinidia*. Manual curation, including reannotation, was conducted for genes with ambiguous predictions or inconsistent clade or subclade assignments.

### 4.3. Selection analyzes

To investigate evolutionary dynamics within the EXO70 gene family, we analyzed clade-specific phylogenies for signs of diversification. We focused on three main indicators: (**i**) unusually long branch lengths; (**ii**) patterns of gene amplification or loss; and (**iii**) incongruence with established phylogenetic relationships among *Actinidia* species. Protein alignments were re-examined to deduce the likely reason for long branch lengths and eliminate more trivial genome annotation differences (e.g. missing or incorrectly deduced open reading frames and reading frame changes after short deletions, insertions or variation in intron splice sites). Patterns of potential gene loss were tested by tBLASTn searches of WGS *Actinidia* genome dataset in NCBI to identify likely missed gene annotations. Phylogenetic incongruence was identified by comparing EXO70 gene trees with known *Actinidia* phylogenies [27] to detect branching patterns that are inconsistent with the deduced relationships of the relevant species based on genome-wide phylogenetic construction.

After selecting the EXO70E1 subclade as one of particular interest (see above), a more detailed analysis of this subclade was conducted using the Adaptive branch-site random effects likelihood (aBSREL) approach [30] to search for episodic diversifying selection. The cDNA sequences corresponding to the four EXO70E1 proteins identified in the reference genome *A. chinensis* var. *chinensis* Red5 were used to search the WGS genome dataset in NCBI to identify *Actinidia* genome sequences for matches that were subsequently aligned (sub-clade by sub-clade) using translation alignment in Geneious. This analysis identified data from three genomes from *A. chinensis* var. *chinensis*, one from *A. chinensis* var. d*eliciosa,* two genomes from *A. eriantha,* two genomes from *A. rufa*, and one genome each from *A. polygama*, *A. melanandra*, and *A. arguta.* The sub-clade alignments were then combined into the complete EXO70E1 clade alignment and manually corrected to conform with the most parsimonious diversification trajectory between the four genes in this clade. Sequence data after indels that affected reading frames were deleted to reduce the effects of random sequence drift after gene inactivation events and stop codons removed to allow the data to be analyzed with aBSREL (v2.5) on Datamonkey 2.0 [31,51]. This analysis uses an adaptive random effects branch-site model framework to test if branches have evolved under positive selection, an evidence ratio threshold of 100 was used to identify possible nodes and/or genes showing signs of episodic selection. A branch tested tree depicting support for selection was constructed using the General Time Reversible (GTR) model for nucleotide substitution. Data collected includes ω values (the dN/dS or non-synonymous to synonymous substitution ratio) and empirical Bayes factors for the branches/nodes, genes and sites under potential selection. ConSurf analysis [28,29] was performed on the *Ath*EXO70B1 protein to identify the regions that show the highest degree of conservation at the surface of the protein.

### 4.4. Yeast Two-Hybrid Assays

Selected *AccEXO70* clades were synthesized and cloned into the pTwist Gateway entry vector (Twist Bioscience) and recombined into the prey vector pDEST22. Full-length *Acc*RIN4_1 was amplified from kiwifruit cDNA and cloned into the bait vector pDEST32. Plasmid pairs were co-transformed into the *Saccharomyces cerevisiae* strain MaV203 following the ProQuest™ Y2H protocol (Invitrogen). Transformants were selected on synthetic dropout (SD) medium lacking Leu and Trp (SD/-Leu/-Trp) and tested for interaction on SD/-Leu/-Trp/-His and SD/-Leu/-Trp/-His/-Ade. Activation of the LacZ reporter gene was visualised by blue coloration on SD/-Leu/-Trp plates containing X-gal. Strong, weak and non-interacting controls from the ProQuest system were used to verify assay performance.

### 4.5. Western blot analysis and co-immunoprecipitation

*Acc*EXO70B1 and *Acc*RIN4_1 from kiwifruit were cloned into pGWB15 and pGWB12, respectively, under the control of the 35S promoter. *N. benthamiana* leaves were syringe-injected using Agrobacterium (GV3101) mixture harboring HA:*Acc*EXO70B1 and FLAG:*Acc*RIN4_1 in equal amounts (OD_600_=0.4 each). Infiltrated tissues (0.5 g per sample) were collected 2 days after infiltration and proteins were extracted with 1 mL of Pierce^TM^ IP Lysis Buffer (87787, Thermo Scientific^TM^). Extracted protein were immunoprecipitated using Pierce™ Anti-HA Magnetic Beads (88836, Thermo Scientific^TM^). Precipitated proteins were run on a 4–12% SDS-PAGE gel. Western blots were probed using HRP-conjugated antibodies in PBS containing 0.2% I-Block (T2015, Invitrogen) and 0.05% Tween 20. Detection was achieved using Clarity Max Western ECL (1705060S, Bio-Rad). Antibodies used were F1804 (anti-FLAG, Sigma-Aldrich), A8592 (anti-FLAG-HRP, Sigma-Aldrich), H9658 (anti-HA, Sigma-Aldrich), 12013819001 (3F10, anti-HA-HRP, Roche, Basel, Switzerland).

## 5. Conclusion

This study presents the first comprehensive characterization of the EXO70 gene family in kiwifruit (*Actinidia* spp.), combining genome-wide identification, evolutionary analysis, and functional validation. We identified 217 EXO70 genes across five *Actinidia* species and classified them into nine major clades consistent with known plant EXO70 groupings. Phylogenetic and synteny analyzes revealed clade-specific expansions and diversification events, particularly within the EXO70E and EXO70H clades.

Evidence of diversifying selection—through gene loss, duplication, and sequence divergence—was strongest in the EXO70E1 and EXO70H4 subclades, suggesting adaptive evolution linked to biotic stress. Functional assays confirmed that two EXO70 clades, *Acc*EXO70B1 and *Acc*EXO70E2, physically interact with the immune regulator RIN4, extending the model of EXO70-mediated immunity from *Arabidopsis* and rice to a perennial fruit crop.

These findings support the hypothesis that specific EXO70 proteins act as hubs integrating vesicle trafficking and immune signaling. The lineage-specific diversification of EXO70 genes in *Actinidia* suggests that these proteins may be important components of kiwifruit’s immunity toolkit.

Together, this work provides a genomic and functional framework for investigating EXO70 roles in plant immunity and lays the foundation for future research aimed at improving disease resistance in kiwifruit. Targeted manipulation of EXO70 genes through molecular breeding or genome editing may offer promising strategies for developing more resilient cultivars.

## Declarations

Ethics Approval and Consent to Participate: Not applicable.

Consent for Publication: Not applicable.

Availability of Data and Materials: Available upon request or in supplementary files.

Competing Interests: The authors declare no competing interests.

## Authors’ Contributions

Wei Cui conceived and led the project, secured funding, performed the yeast two-hybrid (Y2H) analysis, and drafted the original manuscript; Cecilia Deng carried out the phylogenetic analyzes; Minsoo Yoon conducted the co-immunoprecipitation (co-IP) experiments; Erik Rikkerink performed in-depth analyzes of the phylogenetic trees and selection pressures and contributed expertise to the discussion on the evolutionary dynamics of the gene family; Viktor Žárský contributed the conceptual framework for the manuscript and provided insights into the functional diversity of the EXO70 protein family.

## Acknowledgements

We thank the kiwifruit genomics community for providing access to genomic resources. We also acknowledge Kim Snowden, Joanna Bowen, Mark Andersen and Robin MacDiarmid for their valuable comments and proofreading of the manuscript.

## Supplementary Information

Supplementary materials including sequence datasets, alignments, and phylogenetic trees are available in Additional Supplementary Sequence Files SS1-3:

**SS1:** EXO70 peptide sequences in plant genomes (kiwifruit ‘Hongyang’ v3, *Arabidopsis*, rice and tomato)

**SS2:** Alignments of plant EXO70 sequences

**SS3:** EXO70 peptide sequences in kiwifruit genomes including *A. arguta, A. chinensis Red5,* A. *chinensis HongYang, A. eriantha, A. melanandra*, and *Arabidopsis*.

### 1. Supplementary Methods

#### 1.1. Identification of Loci with Long Branch Lengths

The rationale for identifying long branch lengths in phylogenetic trees typically signal a higher-than-expected rate of sequence divergence among otherwise closely related sequences. This pattern can indicate either rapid evolutionary divergence or, in cases where genomes are incomplete, disparities in genome sequencing quality and/or annotation accuracy. For example, annotation artifacts—particularly errors in open reading frame (ORF) prediction— can result in spurious divergence. A bias toward long-branch loci from specific species or subsets, along with the prevalence of truncated or incomplete proteins, often points to such annotation-related issues.

#### 1.2. Identification of Amplified Loci

The detection of gene amplification across genomes of differing quality and annotation strategies presents unique challenges. Partial or fragmented gene calls may inflate locus counts by splitting a single gene into multiple entries. To mitigate this, we excluded proteins covering less than 50% of the conserved EXO70 core from gene counts. Furthermore, recent genome annotations often include haplotype-specific alleles, unlike earlier annotations that presented consensus sequences. For example, the *A. chinensis* Red5V2 genome distinguishes primary from haplotype variant alleles, and only primary alleles were included in our analyzes. In contrast, the *A. melanandra* genome includes haplotype alleles without such differentiation, complicating the identification of true gene amplifications.

#### 1.3. Identification of Subclades Showing Phylogenetic Incongruence

To identify subclades with unexpected branching patterns, we compared our phylogenetic trees to known species relationships outlined by Liu et al. [27]. For instance, alleles from the two *A. chinensis* genomes should cluster more closely with each other than with orthologs from other species. Species from the “hairy or spotted fruit” clade (*A. chinensis*, *A. eriantha*, and *A. rufa*) are expected to diverge from the smooth-skinned fruit group (*A. polygama*, *A. melanandra*, and *A. arguta*). Deviations from this pattern suggest phylogenetic incongruence, possibly indicative of selection.

### 2. Analysis of Specific EXO70 Clades

Results for the EXO70E and EXO70H clades are discussed in the main text. The phylogenetic trees for the other EXO70 clades were examined but did not yield convincing signatures of selection, where missing, incongruent or long branches were identified they were due to poor annotation, sequence quality and/or genes in the process of degenerating. Results for analyzes are discussed under their respective phylogenetic trees below. The long branches for many of the Rosid *Arabidopsis* reference proteins are a reflection of their much greater taxonomic distance from the other proteins in these trees derived from the Asterid genus *Actinidia*.

### 3. Legends for Supplementary Figures

Clade-specific phylogenetic tree of EXO70 proteins, using the same identifying colors as shown in Figure 2A for Figures S1 to S7.

**Figure S1.**
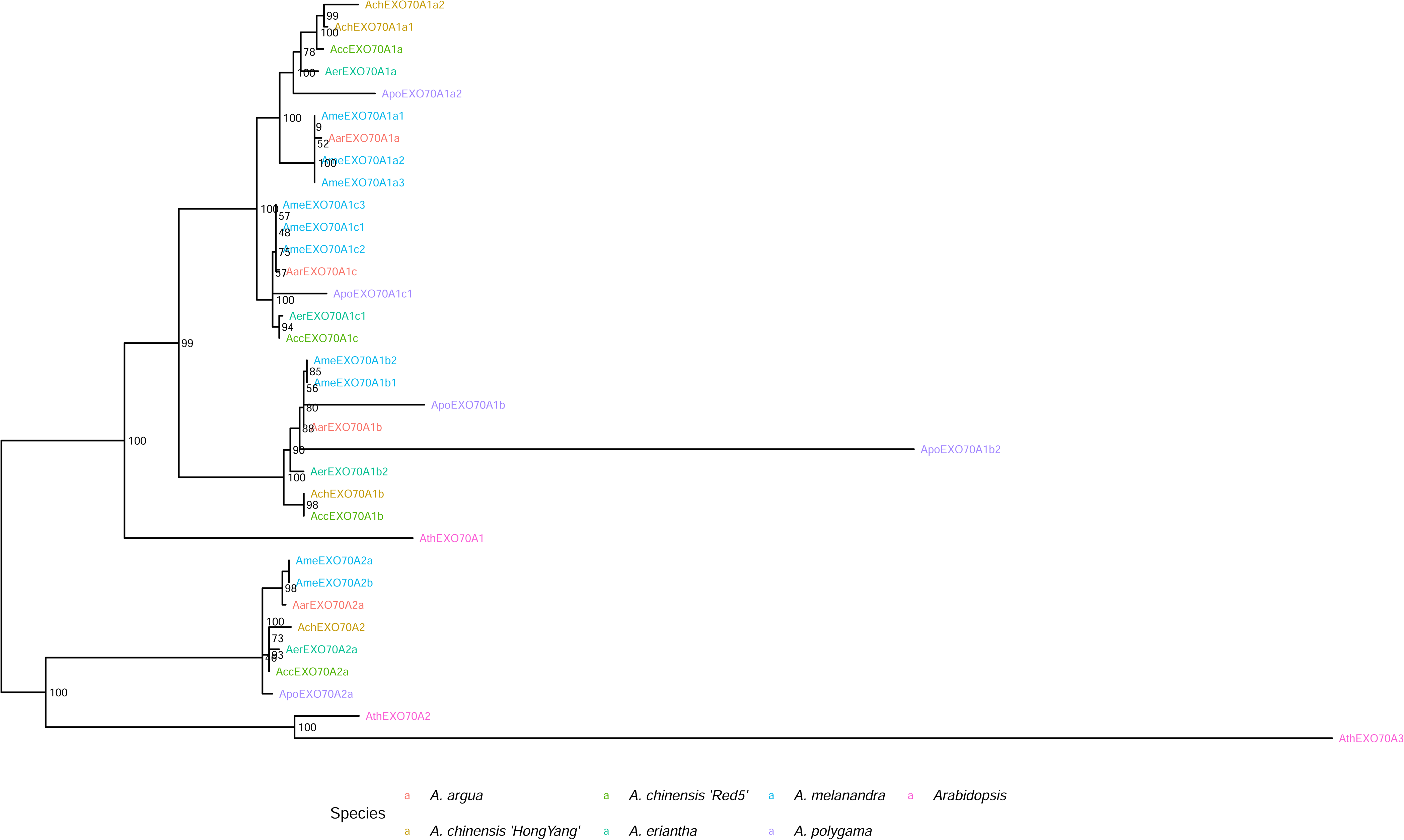
***Apo*EXO70A1b2:** Long branch length due to fusion of the first 5 exons with an exon that has no homology to other EXO70 proteins. Likely due to erroneous gene call and/or a C-terminal deletion/fusion event.

**Figure S2.**
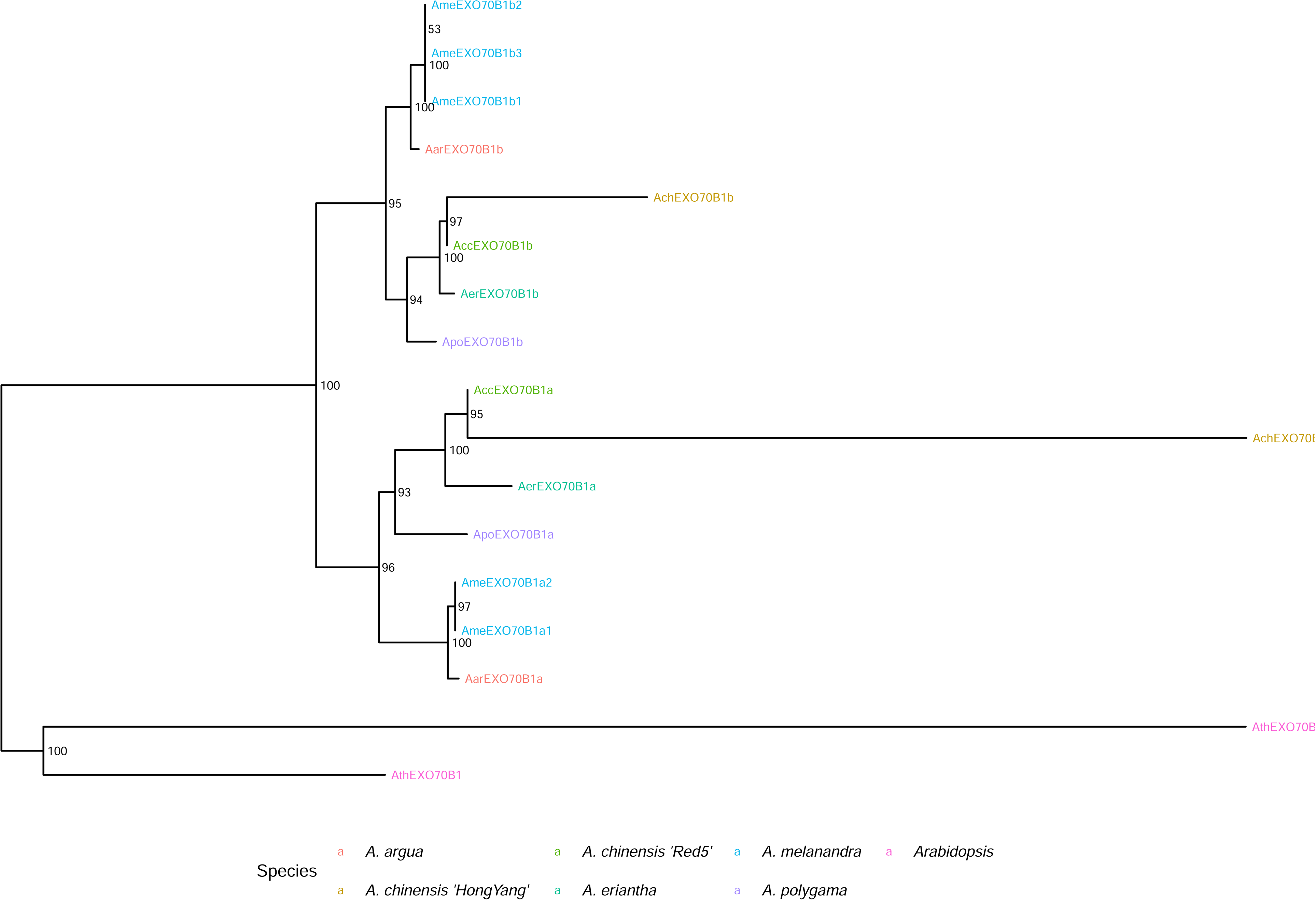
***Ach*EXO70B1 Long Branches**: *Ach*EXO70B1b and *Ach*EXO70B1c exhibit unusually long branches, but both annotations are likely flawed due to missing coding segments. tBlastN analysis revealed near-identical alleles in the Hong Yang genome, suggesting annotation errors. ***At*EXO70B2:** The long branch of *At*EXO70B2 likely reflects true divergence, the EXO70B2 clade appears unique to Brassicaceae.

**Figure S3.**
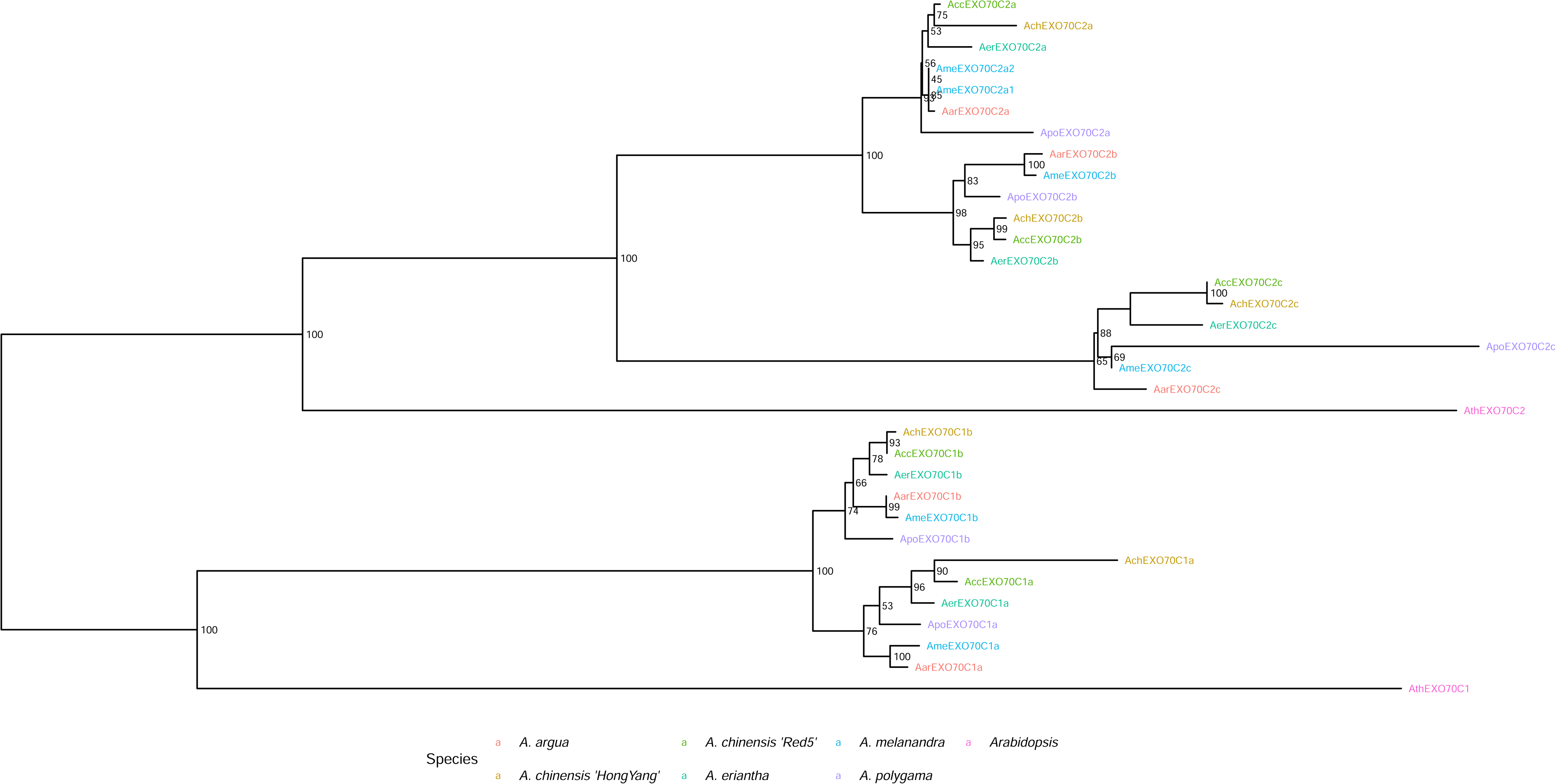
***Ach*EXO70C1a**: Long branch length due to duplication event of leading to a partial C-terminal gene fragment of EXO70C1a being predicted. Further tBlastN analysis indicates that there are a number of frame shifts and/or stop codons in the N -terminal part of the AchEXO70c1a sequence indicating this duplicated copy is in the process of degenerating. There is another potential AchEXO70C1a locus in this genome which may have been missed by the annotation. ***Apo*EXO70C2c**: Long branch length due to only a partial gene fragment based on two exons being predicted. Further tBlastN analysis indicates that other exons do have matching sequences but that this gene is probably not complete in *A. polygama*.

**Figure S4.**
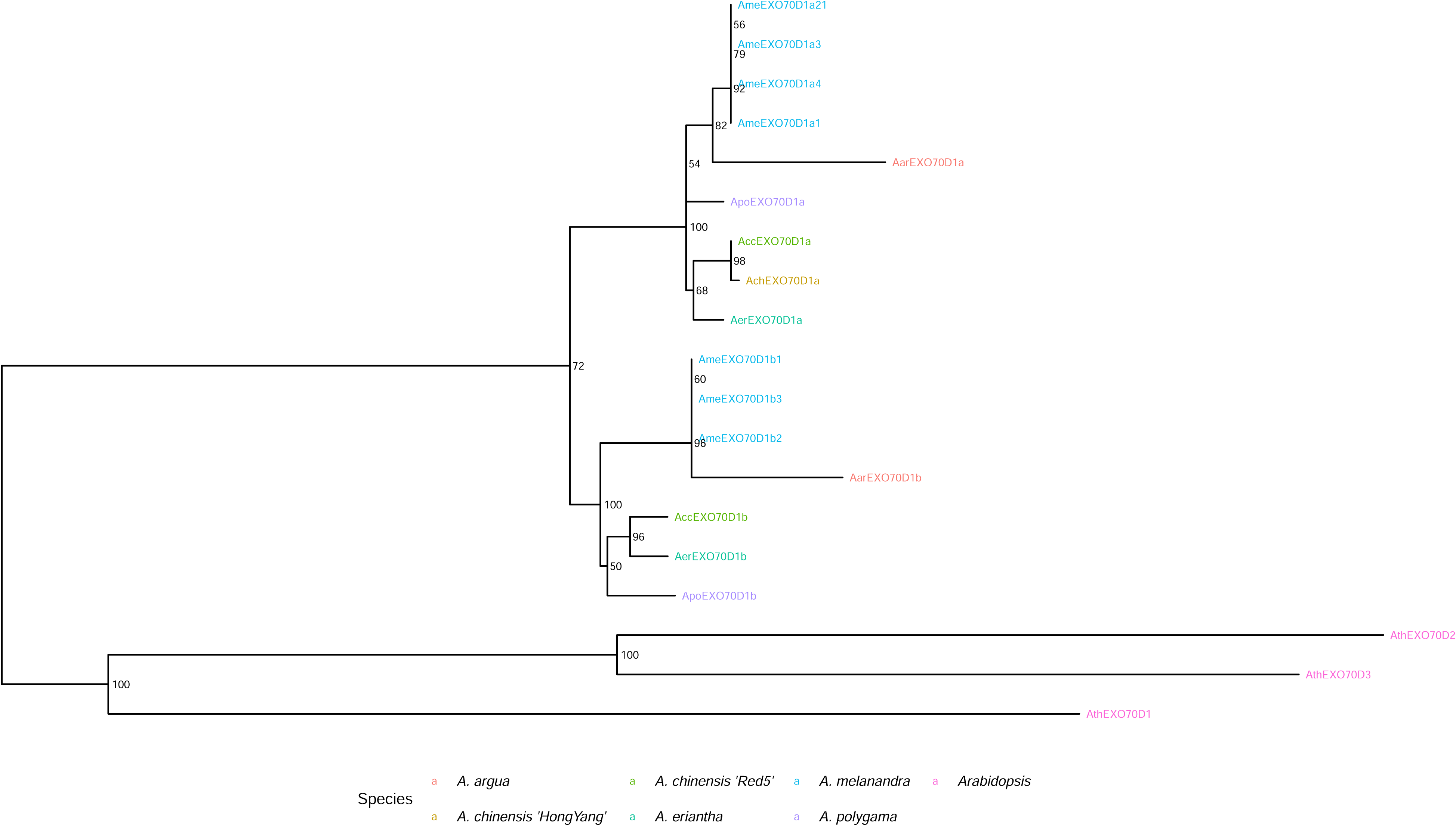
There were no obvious signs of selection in any of the subclades of this tree.

**Figure S5.**
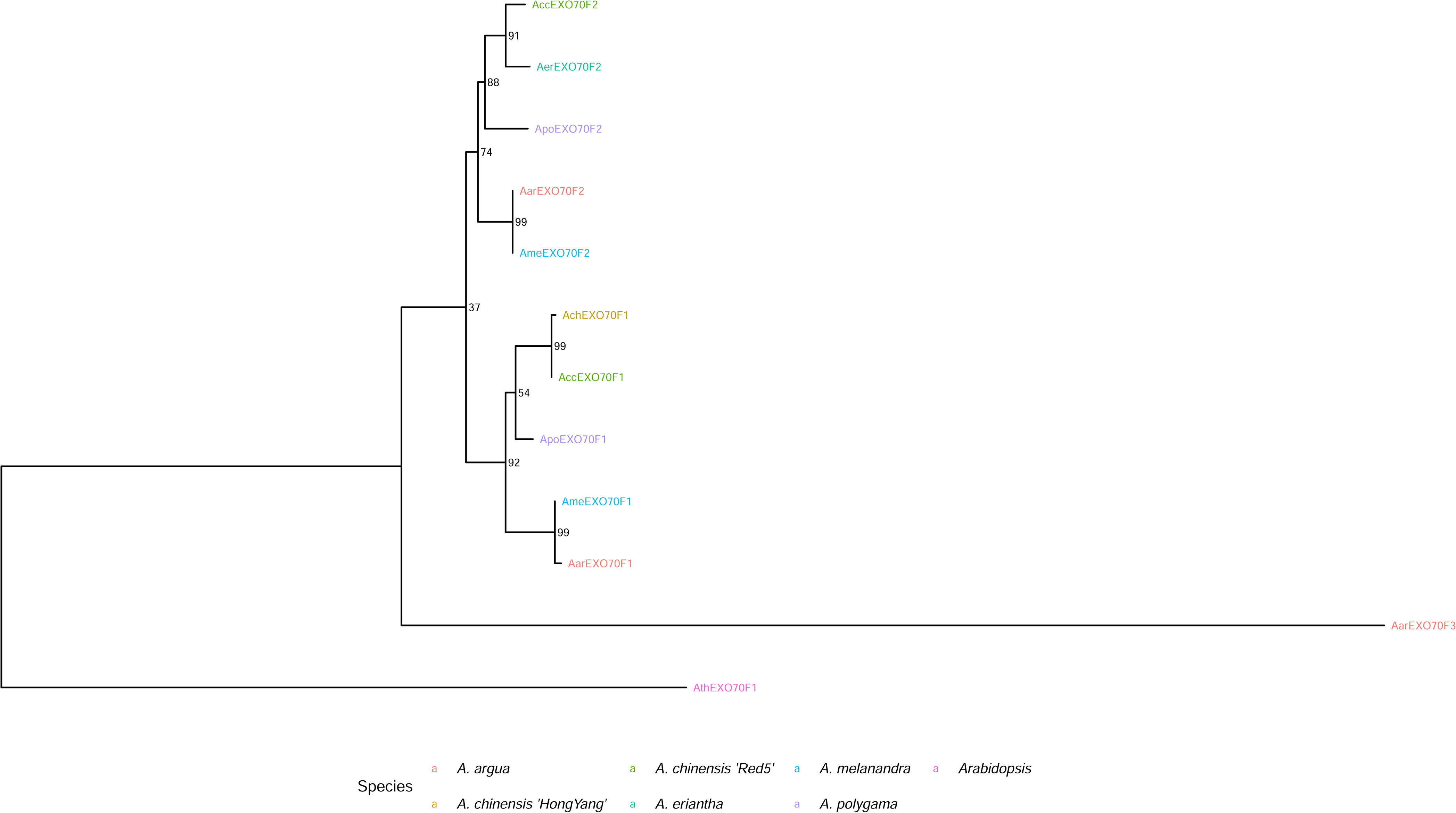
***Aar*EXO70F3**: The long branch of this gene and its unique presence in *A. arguta* suggest past amplification. However, frameshifts in this sequence imply degeneration rather than diversification may account for this long branch instead.

**Figure S6.**
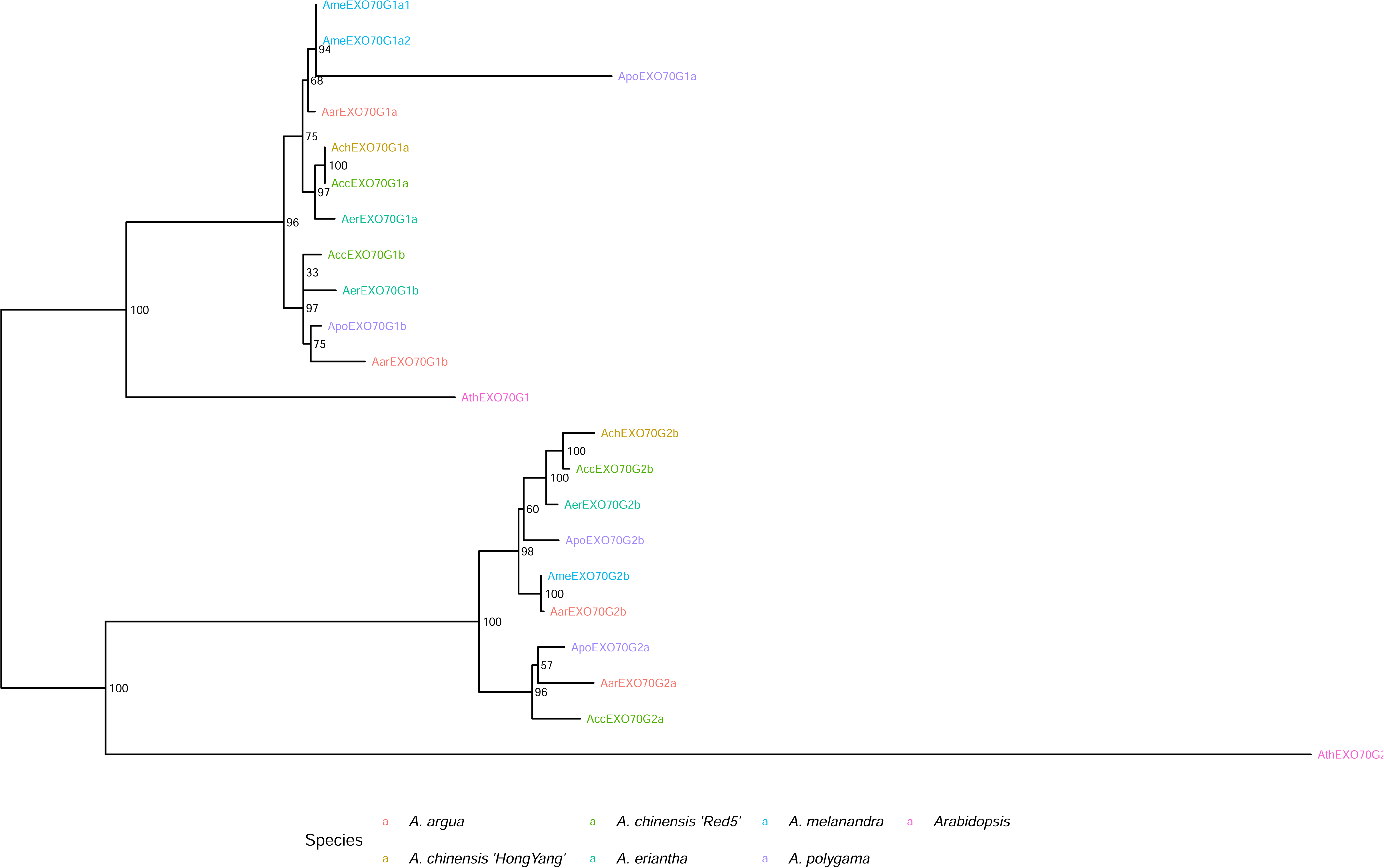
***Apo*EXO70G1a**: Long branch length caused by a partial gene being called that has missing gene fragments. This was probably caused by a number of different frameshifts and stop codons present interfering with translation of this gene i.e. the gene sequence is poor or the gene is in the process of degenerating.

**Figure S7.**
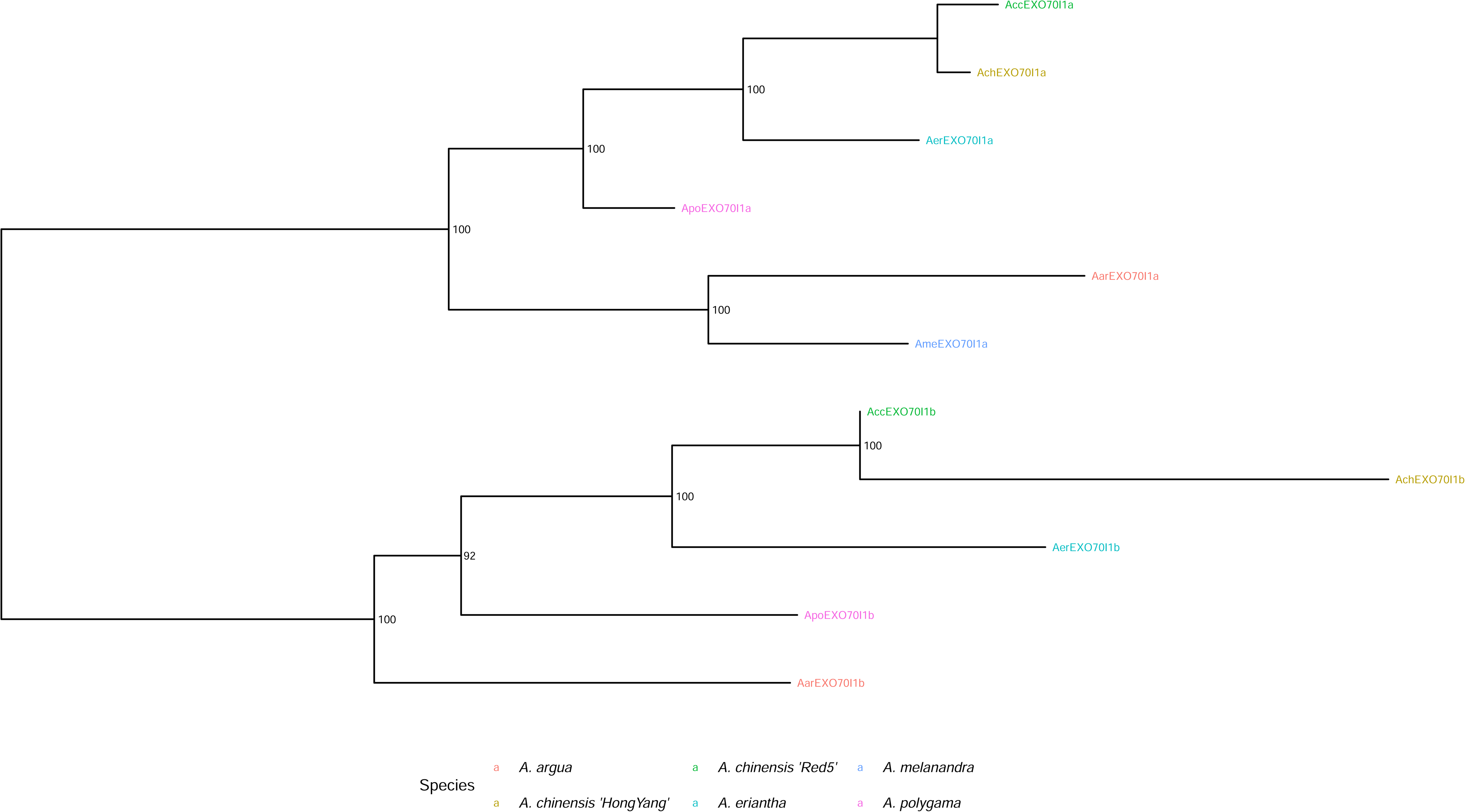
***Ach*EXO70Ib**: The long branch length of this gene is caused by incorrect annotation of this gene leading to a missing exon at the N-terminus as well as additional translated sequence at the C-terminus that does not match other EXO70Ib alleles. The tBlastN analysis suggests these are either annotation errors as matching sequences are present for each of the exons or that some of the intron splice sites have been compromised by sequence variation.

**Figure S8.**
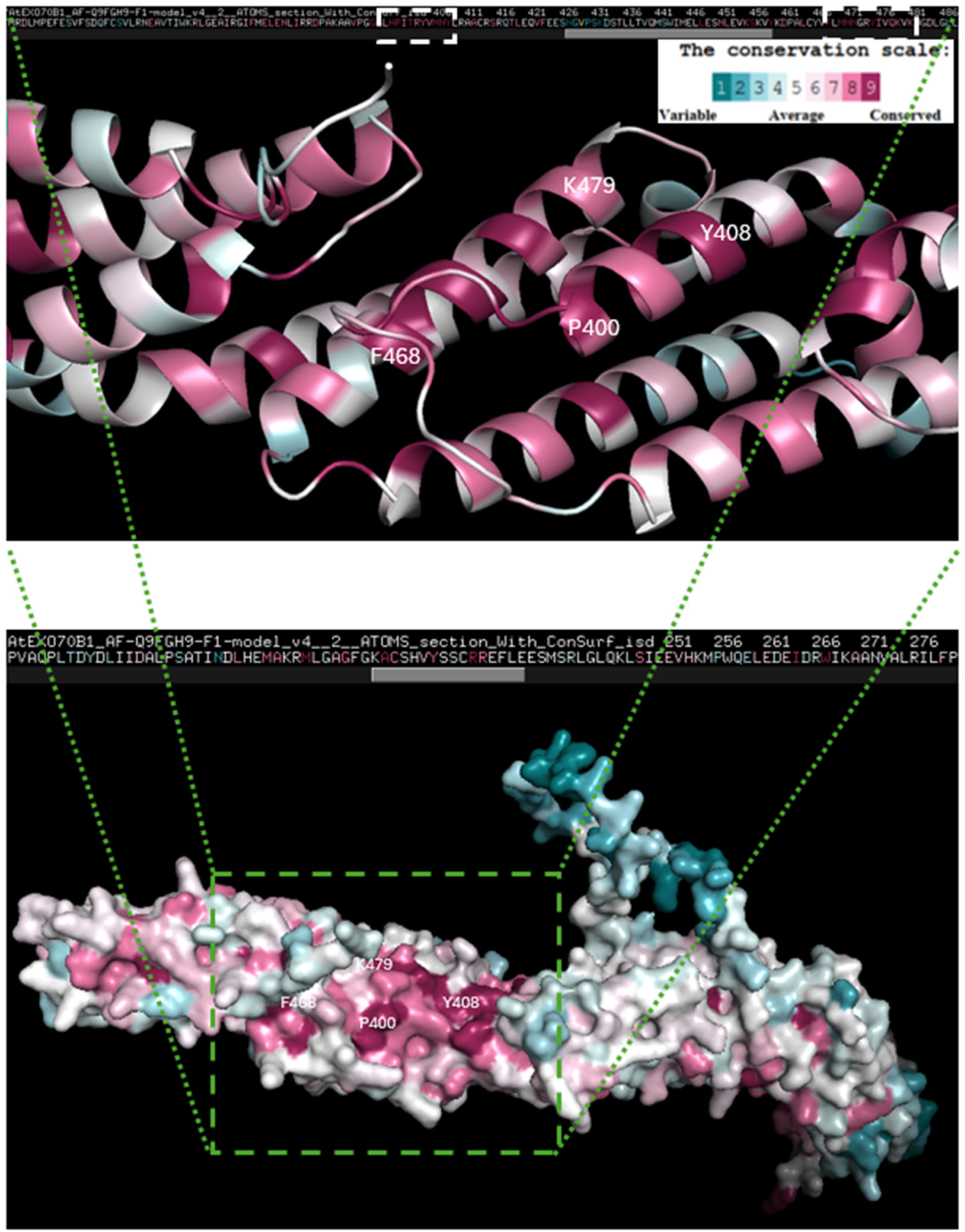
PyMol depiction of the structure of *Ath*EXO70B1 analyzed by Consurf to identify the degree of conservation of residues at the surface of the protein. The two most highly conserved alpha helices in the middle of EXO70B1 are identified in the bottom structure by the dark red shade and numbering of the corresponding start and end residues of the helices. A cartoon-style magnification of this region is presented at the top with the conservation scale. Matching protein sequences for these helices are surrounded by a white dashed boxes in the protein sequence presented above the top magnification.

**Figure S9.**
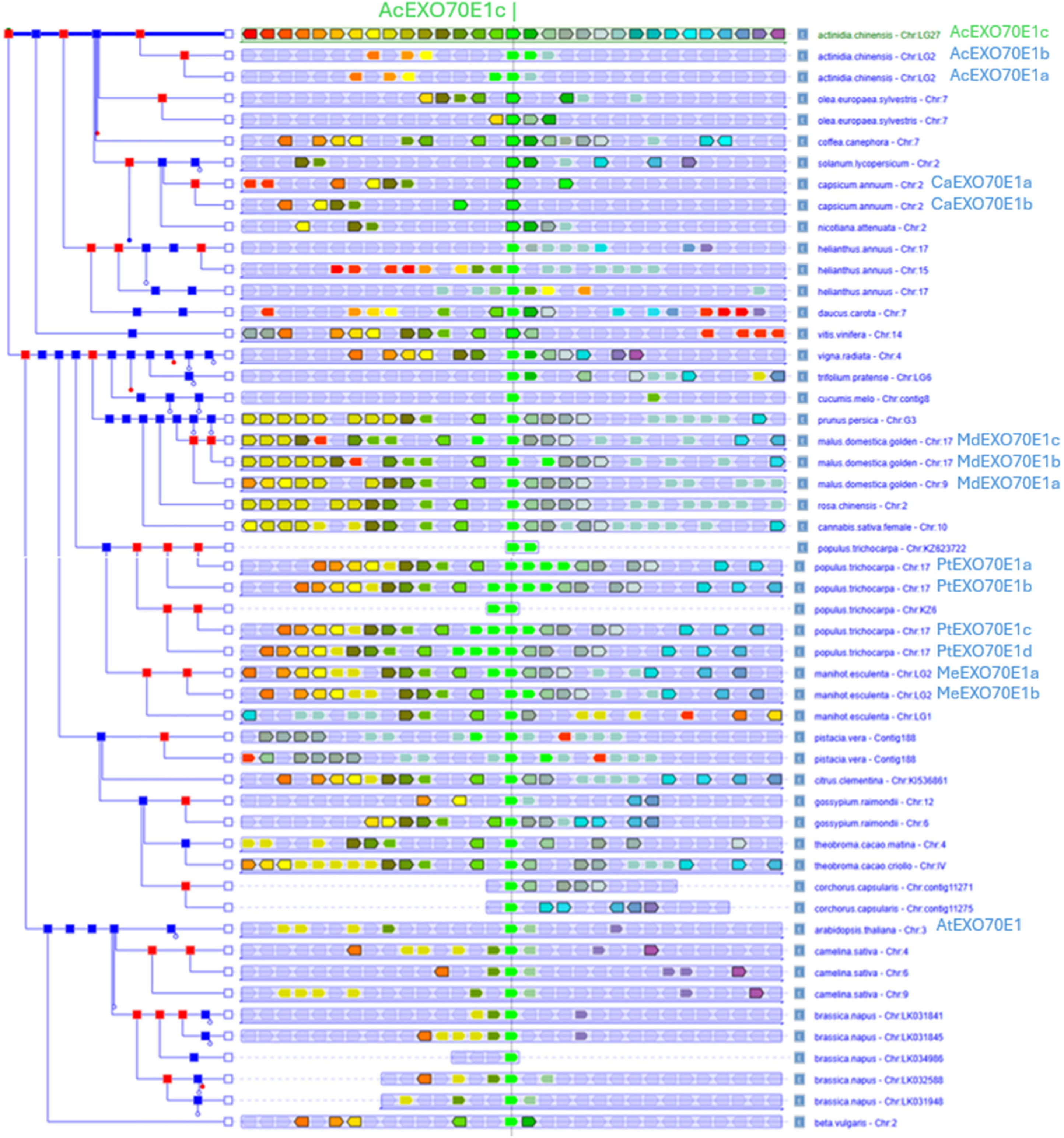
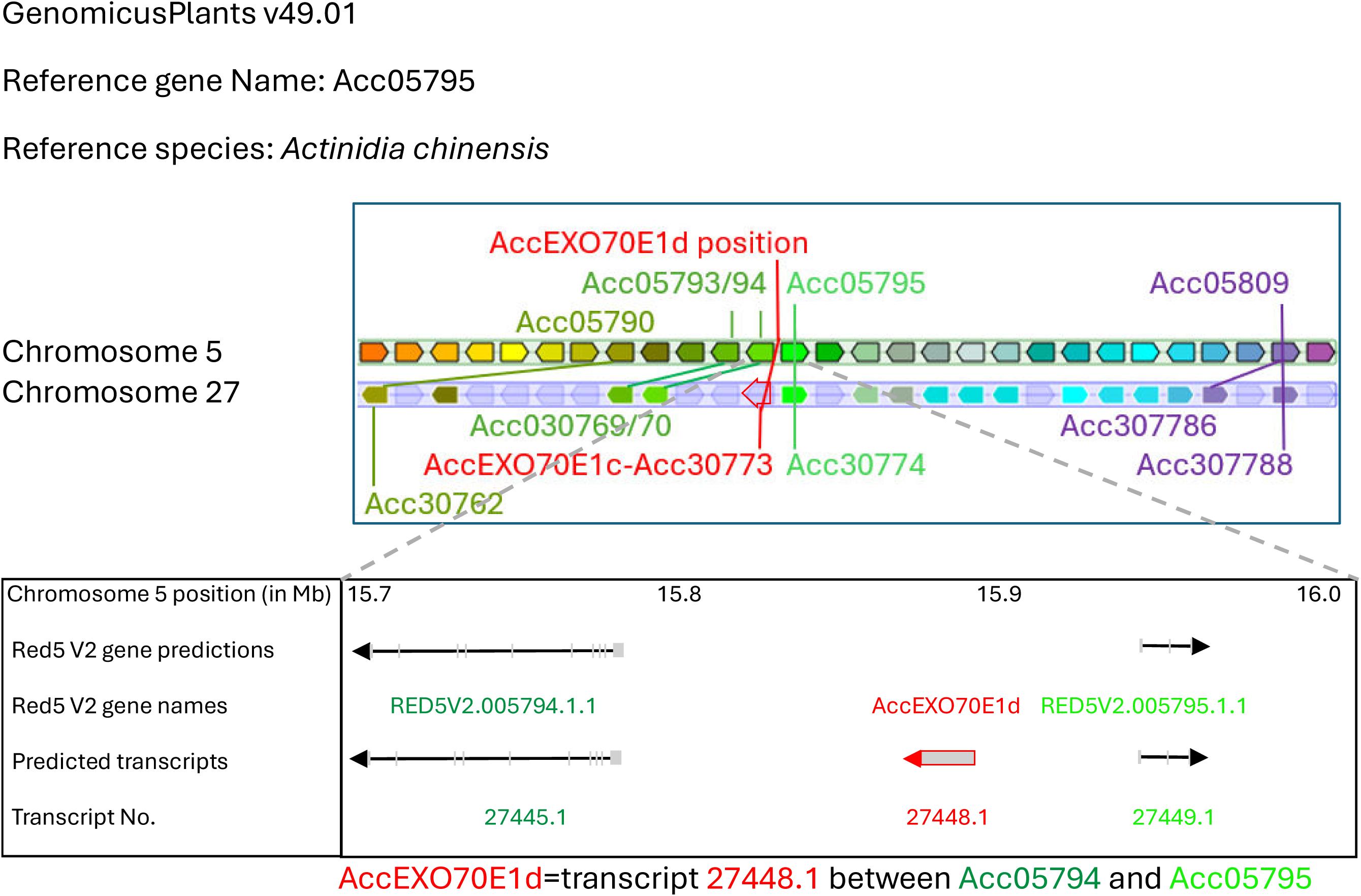
**A**) Evidence of microsynteny around the kiwifruit EXO70E1a, b and c loci when compared to other plant genomes from asterid and rosid species present in EnsemblePlants. In this Genomicus Plants summary orthologous genes have the same shade of color between species and indicate retention of micro-syntenic regions between species. Multiple light green arrows (e.g. on Chr17 in *Malus* and *Populus*) indicate localized amplifications of EXO70E loci on these chromosomes. **B**) Identification of a new EXO70E1 locus (*Acc*EXO70E1d) by using the flanking predicted genes in GenomicusPlants that contains *A. chinensis* Red5 Version 1 gene predictions (top panel). Genes homoeologous to the genes flanking *Acc*EXO70E1c on chromosome 27 were used to identify an unannotated transcript (27448.1) on chromosome 5 (bottom panel) that is highly similar to *Acc*EXO70E1c and is present in all other *Actinidia* genomes and therefore indicates the presence of an extra unannotated EXO70E1 locus EXO70E1d. Locus initially identified in the genome of *A. chinensis* var. *chinensis* Red5.

**Figure S10.**
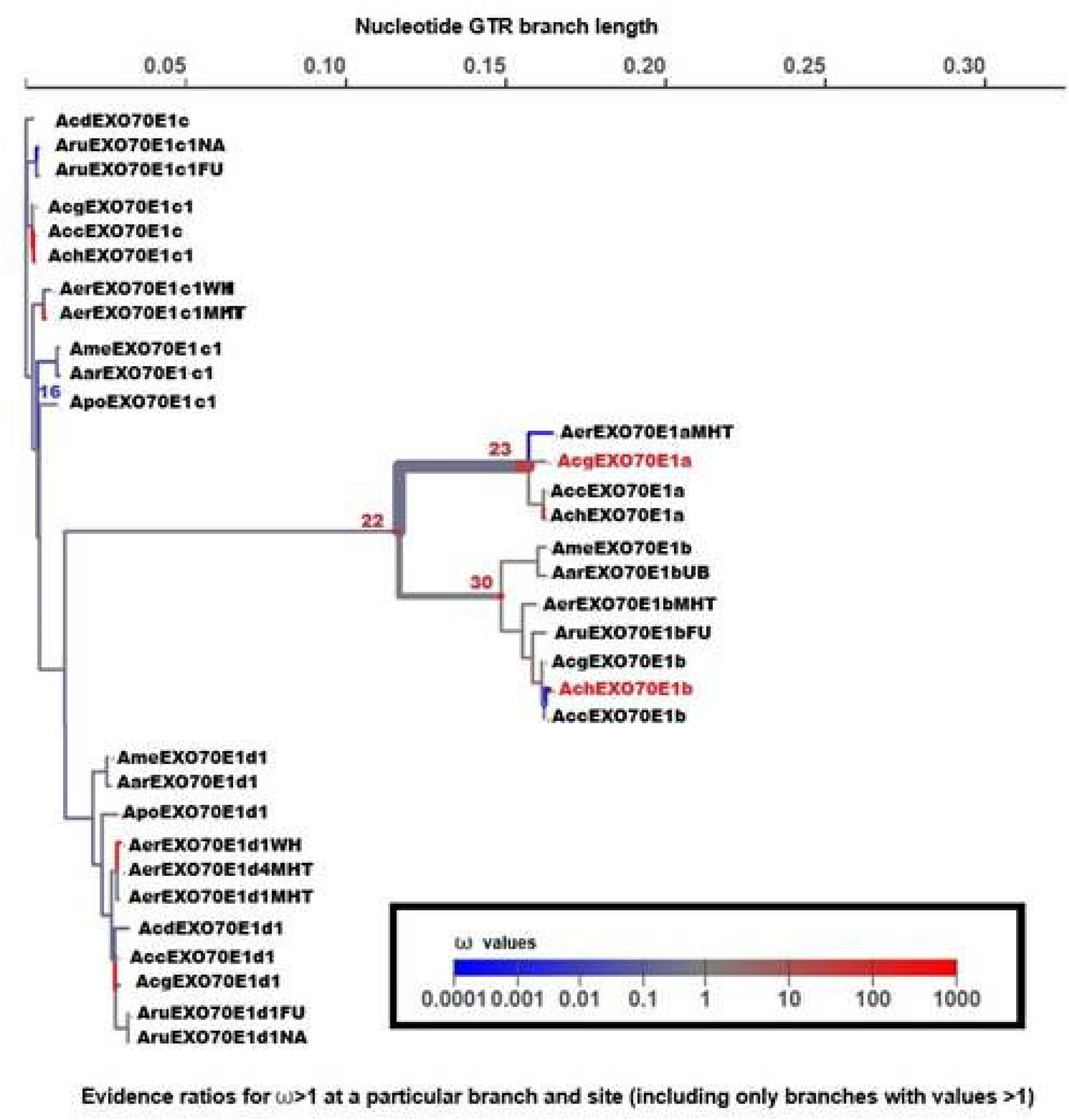
Branch testing for selection mapped onto tree from translation based alignment of the kiwifruit EXO70E1 clade using aBSREL analysis. Branches/genes with ω values >1 are indicated and colored according to the given ω scale. Selected values are given in the Table S2. Gene names identify the genome of origin of the relevant cDNA sequence as follows: Acc *Actinidia chinensis* var. *chinensis* Red5; Ach *Actinidia chinensis* var. *chinensis* Hongyang; Acg *Actinidia chinensis* var. *chinensis* Guimi No. 2; Acd *Actinidia chinensis* var. *deliciosa* Isolate 284548; Aer *Actinidia eriantha* varieties WH=White MHT= MHT-2021; Aru *Actinidia rufa* varieties FU=Fuchu, NA=Nakamura B; Apo *Actinidia polygama* Hokkaido-male; Ame *Actinidia melanandra* ME02_01; Aar *Actinidia arguta* var. h*ypoleuca* Ubaishi-male.

### 4. Supplementary Tables

**Table S1:** Summary of potential diversifying selection in the EXO70E1 clade.

|  | EXO70E1 clade |  |  |  |
| --- | --- | --- | --- | --- |
| ***Actinidia* species** | EXO70E1a | EXO70E1b | EXO70E1c | EXO70E1d |
| *A. chinensis* Red5 | N | P | P | P |
| *A. chinensis* Hongyang | P | P | P | P |
| *A. eriantha* | N | N | P | P |
| *A. polygama* | A | N | P | P |
| *A. melanandra* | N | D | P | P |
| *A. arguta* | N | D | P | D |

A table summary highlighting the differences between the four EXO70E1 loci in kiwifruit. **P**=present, **A**=absent, **N**=non-functional/truncated, **D**=divergent protein sequence. Presence, absence or truncation was determined by tBlastN searches of Red5 genes against *Actinidia* genomes available in the NCBI WGS database.

**Table S2:**
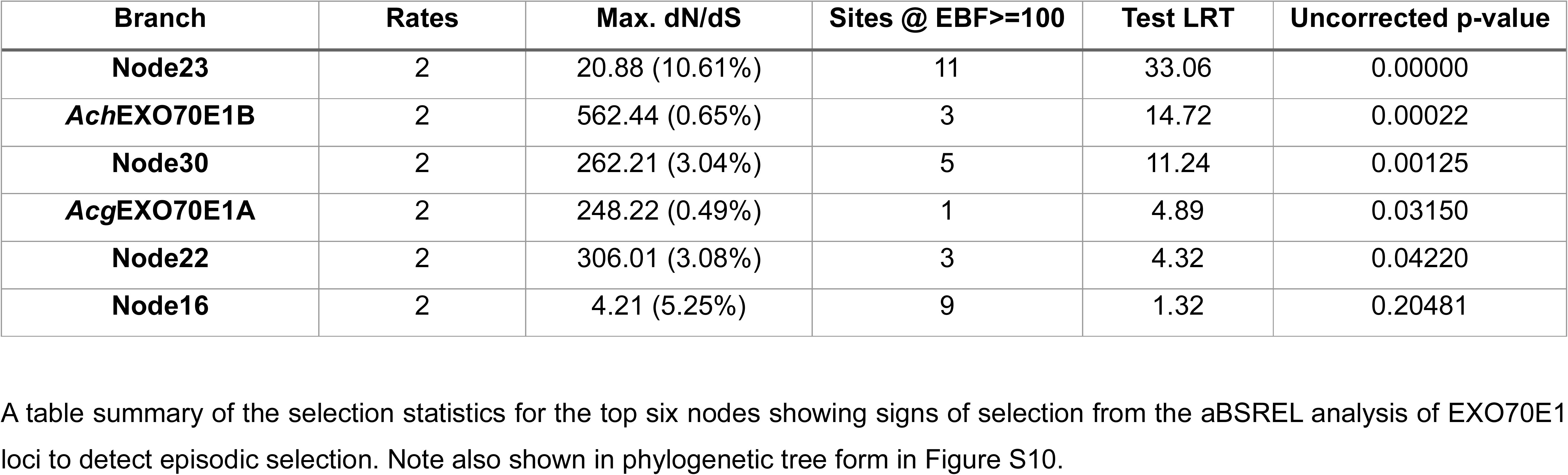
Summary of kiwifruit EXO70E1 nodes/genes showing most significant selection.

